# VoroIF-GNN: Voronoi tessellation-derived protein-protein interface assessment using a graph neural network

**DOI:** 10.1101/2023.04.19.537507

**Authors:** Kliment Olechnovič, Česlovas Venclovas

## Abstract

We present VoroIF-GNN, a novel single-model method for assessing inter-subunit interfaces in protein-protein complexes. Given a multimeric protein structural model, we derive interface contacts from the Voronoi tessellation of atomic balls, construct a graph of those contacts, and predict accuracy of every contact using an attention-based graph neural network. The contact-level predictions are then summarized to produce whole interface-level scores. VoroIF-GNN was blindly tested for its ability to estimate accuracy of protein complexes during CASP15 and showed strong performance in selecting the best multimeric model out of many. The method implementation is freely available at https://klimentolechnovic.github.io/voronota/expansion_js/.

## 1 Introduction

The breakthrough in protein structure prediction, brought about by AlphaFold [1], provided possibilities to obtain high-accuracy models of single-chain structures for most of the known protein sequences [2, 3]. However, prediction of structures for protein multimers still remains a major challenge [4–6]. In addition to AlphaFold, there are other new methods that are able to predict structures of protein multimers [7–9]. Since structures of protein complexes are key to understanding function, there is no doubt that the active development of new methods aimed at prediction of multimeric structures will continue. Consequently, the ability to select the most accurate multimeric model independently of how it was obtained is becoming of utmost importance. Not surprisingly, the estimation of model accuracy (EMA) for multimeric structures recently received a lot of attention in the field and was introduced as a new prediction category in CASP15.

In our previous studies we found that the proteinprotein interface-focused accuracy assessment might be effective in identification of best multimeric models. For example, in CASP14 our predicted complexes had the overall most accurate protein-protein interfaces [10] despite the fact that the structure accuracy of monomers of our multimeric models was not the best. These results suggested that with the increased subunit accuracy, such as that provided by AlphaFold, the focus on the inter-subunit interface accuracy might be even more rewarding. Based on this assumption, we developed a new single-model method, VoroIF-GNN, for the assessment of protein-protein interaction interfaces in multimeric protein structures. The method is based on the bottom-up approach, that is, initially every inter-chain residue-residue contact is assigned an accuracy score and then those scores are summarized to derive a single global accuracy score for either the whole set or a subset of interfaces in the multimeric structure.

We tested the ability of VoroIF-GNN to make reliable accuracy estimates for multimeric structures in the CASP15 blind-mode setting as the “VoroIF” group. In addition, VoroIF-GNN was a key component of our newly developed multi-model method, VoroIF-jury, which was primarily used for model ranking and selection in protein assembly modeling by our human group (“Venclovas”). For reference, under the name of “VoroMQA-select-2020” we also included our CASP14 EMA method. Since VoroIF-jury is described in an accompanying paper in this Proteins issue [11], here we focus exclusively on VoroIF-GNN. We describe the underlying ideas and their implementation as well as provide a concise description of its performance in the EMA category of CASP15.

## 2 Materials and methods

### 2.1 Representing inter-chain interactions as a graph of contacts — an interface graph

We use our previously developed methodology [12, 13] to define atom-atom contacts as Voronoi faces — surface patches shared by adjacent Voronoi cells in the Voronoi tessellation of atomic balls of van der Waals radii. We constrain the Voronoi faces inside the solvent accessible surface [14]. A residue-residue contact is defined as a set of relevant atom-atom contacts. Similarly, an inter-chain interface is defined as a set of inter-chain atom-atom contacts.

Figure 1a shows an example of Voronoi tessellation-derived inter-chain interface where atom-atom contacts are be colored by their correspondence to residue-residue contacts. The tessellation is used to determine what contacts are adjacent, i.e. sharing at least one Voronoi edge (Voronoi edges are shown as black line segments in Figure 1a). The adjacency information is used to define an interface graph with nodes representing atom-atom contacts (Figure 1b). Graph of atom-atom contacts can be converted to graph of residue-residue contacts (Figure 1c) by merging nodes and connections (edges) according to the atom-to-residue mapping.

**Figure 1:**
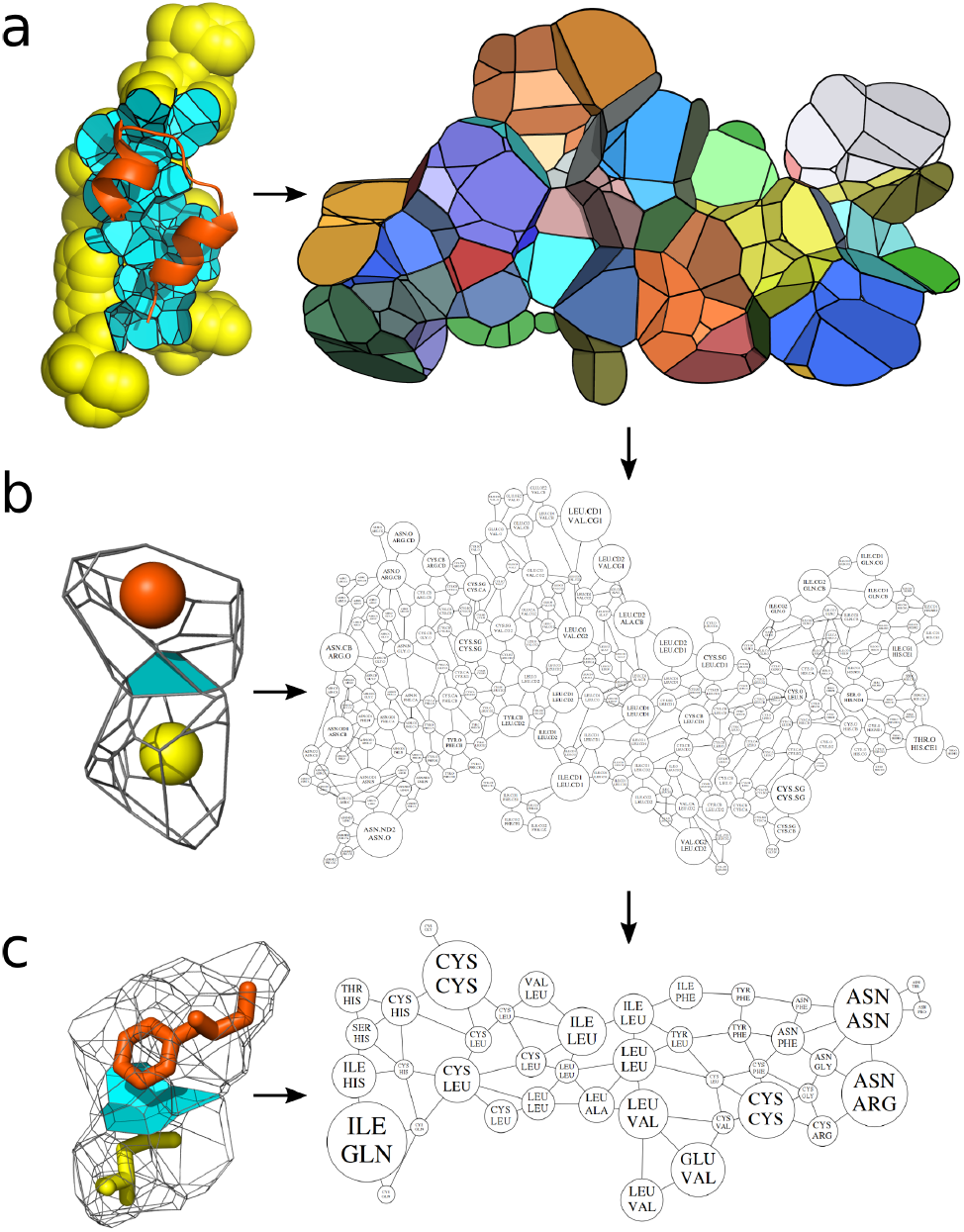
Deriving the interface graph from the Voronoi tessellation of atomic balls: tessellation-derived inter-chain contacts constrained inside the solvent-accessible surface (a); graph of atom-atom contacts (b); graph of residue-residue contacts (c).

### 2.2 Annotating an interface graph with input attributes

We start with a graph of atom-atom contacts and annotate it with computed geometric attributes (Figure 2). For node-to-node edges we set inter-contact border lengths. For nodes we set contact surface areas, contact-solvent border lengths, and sums of intercontact border lengths.

**Figure 2:**
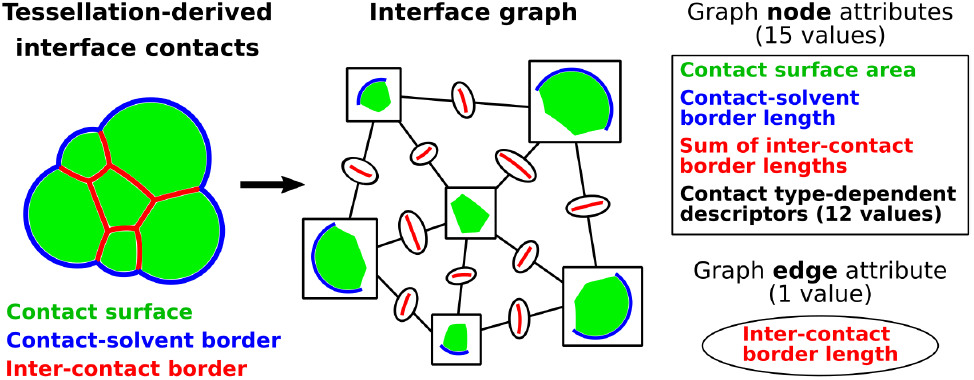
Input attributes of the nodes and edges in an interface graph.

Processing a graph using a graph neural network will require self-connections for every node. For every such connection we set the inter-contact border length value *l* to be equal to the half of the circumference of a disk with the same area *S* as the contact surface area in the node that is being self-connected: 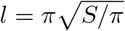.

We also assign contact type-derived attributes to nodes. Every node represents a contact between an atom of type *A* and an atom of type *B*. We take three values from the statistical potential coefficients defined for the VoroMQA [13] method: VoroMQA_(*A,B*)_, VoroMQA_(*A*,solvent)_, VoroMQA_(*B*,solvent)_. This triple of values is unique for every contact type. We also multiply the three potential values by the contact area and obtain three VoroMQA pseudo-energy values — these values represent both the observed situation when atoms of types *A* and *B* are in contact, and an alternative scenario when the atoms are detached from each other and interact with solvent instead. In addition to the six values derived using the proteinspecific VoroMQA potential, we derive analogous six values using the VoroMQA potential adapted for a reduced set of atom types defined by the Knodle [15] atom typing system.

An important trait of most of the described node-level and edge-level attributes is that they are summable (capable of being added). When merging nodes and edges to convert a graph of atom-atom contacts into a graph of residue-residue contacts, we compute residue-level attributes by adding up the relevant atom-level attributes. The only attributes that are not directly summable are VoroMQA potential co-efficients — for residue-residue contacts we replace them with normalized VoroMQA-energy values (the summed VoroMQA-energies divided by the summed areas).

In total, for either atom-atom or residue-residue level interface graphs, we have 1 attribute value for every node-to-node connection (inter-contact border length) and 15 attribute values for every node (3 geometric + 12 VoroMQA-based).

### 2.3 Assigning ground truth values to interface graph nodes

We use the CAD-score method [16, 17] to define a reference-based accuracy estimate for every interchain residue-residue contact. A CAD-score value in [0, 1] interval for a single contact (*i, j*), where *i* and *j* are identifiers of contacting residues, is computed from the contact area *M*_(*i,j*)_ observed in the model structure and the *T*_(*i,j*)_ area observed in the target (reference) structure, as shown in Eq. (1). Examples of modeled interfaces with contacts colored according to CAD-score values are shown in Figure 3.

**Figure 3:**
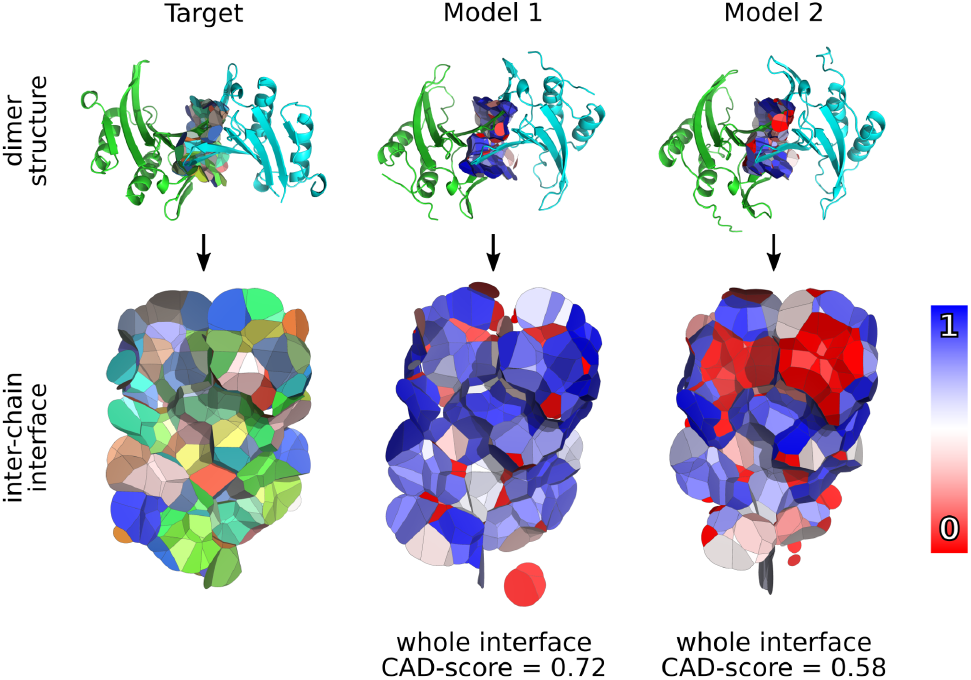
Examples of CAD-score on both global (whole interface) and local (per contact) levels. The interface contact surfaces are colored according to CAD-score values computed for residue-residue contacts.

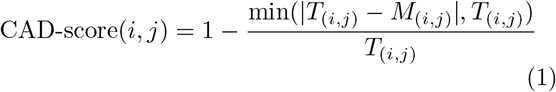

We would like to compute an accuracy estimate for the entire interface as a sum of values for individual contacts. We also would like to use contact areas as weights to reflect uneven contribution of individual contacts to the interface. However, CAD-score values of individual contacts are not summable. Therefore, in Eq. (2) we define the conversion of CAD-score to a pseudo-energy-like value, CAD-goodness. The conversion uses the inverse error function (erf^*−*1^) and can also be defined using the inverse cumulative density function of the normal distribution (qnorm) — this way the conversion can be viewed as converting a probability estimate to a z-score, shifting it by one standard deviation value, and then multiplying the shifted z-score by the contact area. The shift is applied to reduce the penalization of CAD-score values below 0.5.

The CAD-goodness-to-CAD-score transformation is defined in Eq. (3). Plots for both Eq. (2) and Eq. (3) are presented in Figure S1 — they show that CAD-goodness is defined to correlate positively with CAD-score — higher CAD-goodness values correspond to higher CAD-score values. CAD-goodness values can be negative, therefore the total sum of all the interface graph node CAD-goodnesses can be negative, which is expected to happen when the protein-protein complex is modeled grossly incorrectly.

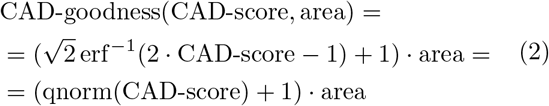

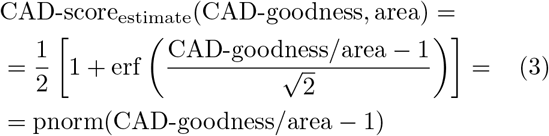

### 2.4 Deriving global interface scores from node-level predictions

We derive a predicted global quality score (called VoroIF-GNN-score) for the whole interface by summing the predicted CAD-goodness values for all the interface graph nodes. We also define normalized-VoroIF-GNN-score — VoroIF-GNN-score divided by the total sum of interface contact areas. Another way to normalize VoroIF-GNN-score is to convert it to predicted CAD-score (pCAD-score) using Eq. (3).

We can compute global scores for any subset of interface contacts. For example, pCAD-score can be computed separately for every residue that has contacts belonging to the inter-chain interface. We also define another whole interface-level score, average-VoroIF-GNN-residue-pCAD-score — an average of pCAD-score values of interface residues.

### 2.5 Measuring performance of predicted global scores

When assessing our scoring approach, we focus on the ability of our global scores to select the best predicted protein-protein complex model out of a set of models of the same target. We evaluate this ability by scoring multiple sets of models and calculating the best model selection rate — how often the selected model (the model with the highest predicted global score, e.g. VoroIF-GNN-score) is the same as the actual best model (the model with the highest reference-based score, e.g. interface CAD-score).

For multiple sets of models, we also calculate the sum of reference-based scores of the selected models. This type of performance evaluation may be more informative in cases when the choice of the actual best model is ambiguous, i.e. when multiple models have similarly high reference-based scores.

### 2.6 Preparing datasets for training, validation and testing

We used rigid body docking to generate protein-protein complex models of different accuracy. We used the database of the PPI3D web server [18] to collect a non-redundant set of 1567 target structures — heterodimeric experimentally-determined assemblies.

We chose to use dimers because we wanted interface graphs that are not overly large — to limit computational resources needed for training. We chose to not use homo-oligomers because we did not want our method to learn to prefer symmetric interfaces. The full set of targets (1567 targets) was randomly split into training (1097), validation (235), and testing (235) sets of targets.

For every target structure, we re-docked its monomers using FTDock [19]. In order to have a more exhaustive set of models, we forced FTDock to not filter docking results using its internal scoring. This way 21440 unique models were generated for every target, 21440 *×* 1567 = 33596480 models in total. Rigidbody docking is known to produce steric clashes — we reduced the number of clashes by rebuilding model side-chains using FASPR [20]. For every model, we computed two reference-based global accuracy scores, the interface CAD-score and the binding-site CAD-score, that differ by their stringency [17]. The interface CAD-score evaluates every residue-residue contact across the interface, whereas the binding site CAD-score only asks whether a residue is part of the interface and does not care whether the individual contacts across the interface are correct. We also computed interface VoroMQA-energy scores and used them to rank the models of every target.

For every ranked list, starting from the model with the best VoroMQA-energy, we selected a sequence of models having interface CAD-score values that differ by 0.1 or more. Then we restricted the ranked list to only the models with interface CAD-score equal to 0 and, starting from the model with the best VoroMQA-energy, selected a sequence of models having binding site CAD-score values that differ by 0.1 or more. As a result, for every target, we got up to 20 models (12.8 models on average). Table 1 summarizes the numbers of selected models for training, validation, and testing. The models cover a wide range of accuracies both in terms of the detailed interface accuracy and the binding-site accuracy (Figure S2abc). Moreover, the selected diverse models, while being mostly incorrect, have better-than-random VoroMQA-energy values and, therefore, can be deceptive decoys for accuracy estimation methods. In our prepared datasets good models and bad models are very often similar in terms of interface size (Figure S2d) and normalized (per unit of area) interface VoroMQA-energy (Figure S2e).

**Table 1:** Summary of targets and docking models used for training, validation, and testing.

| Dataset | Targets | Models |
| --- | --- | --- |
| training | 1097 | 14400 |
| validation | 235 | 2845 |
| testing | 235 | 2814 |

A target structure can be viewed as an ideal model (its both interface and binding site CAD-score values are equal to 1), but we only included such models for training. Validation and testing data sets were deprived of ideal models to make the task of selecting the best model more difficult. For all the selected models, interface graphs of residue-residue contacts were constructed, graph nodes and node-to-node connections (edges) were annotated with input attributes. CAD-goodness values were computed for graph nodes to serve as ground truth values for prediction.

### 2.7 Graph neural network design, training, and validation

We chose to use a graph neural network (GNN) to predict CAD-goodness values for nodes of a residueresidue-level interface graph. Based on our expertise in protein-protein interactions and some basic understanding of modern GNN capabilities, we formulated several considerations about the desired GNN design:

- node-level predictions must be done considering the graph neighborhood of a node, therefore message passing operators must be employed;
- both node and edge features must be utilized in message passing;
- GNN must be trainable to consider multiple aspects of node-edge-node relationships, therefore a multi-headed attention mechanism must be used in message passing;
- not only immediate neighborhood is important for every node, therefore message passing must be implemented in multiple layers.

Following the above considerations, we defined the GNN architecture shown in Figure 4 and implemented it in PyTorch Geometric [21, 22] framework. We used 4 layers of improved graph attention [23] convolutional operators (GATv2Conv) with 8 attention heads and dropout coefficient of 0.25. Exponential linear units [24] were applied after non-output layers.

**Figure 4:**
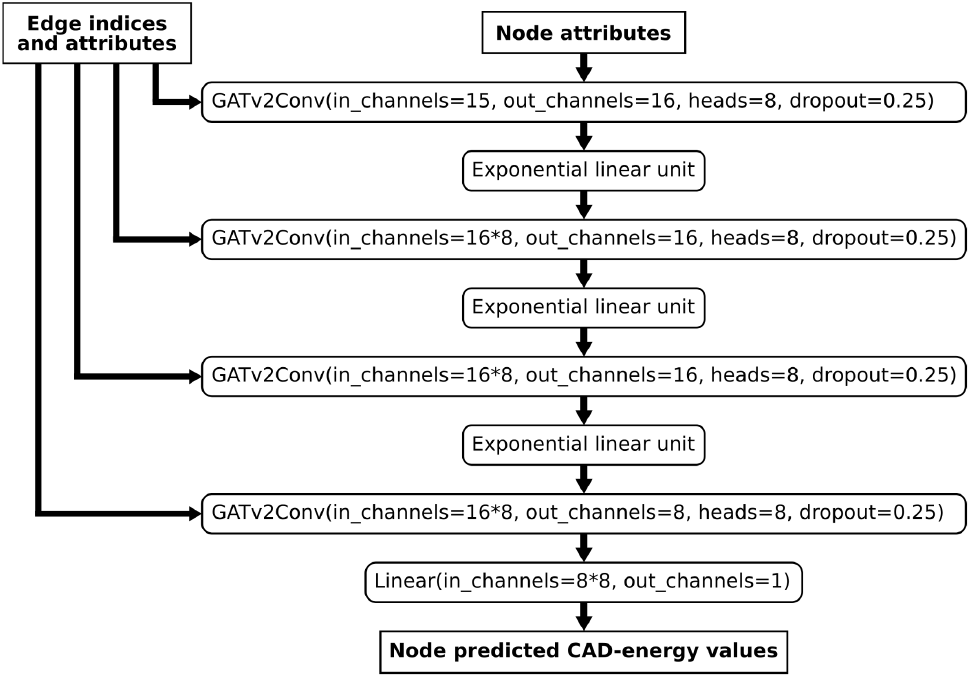
VoroIF-GNN graph neural network architecture.

To train the GNN, we used the training dataset of interface graphs of docking models (described in the previous subsection). Before training, we standardized the data by converting all the input feature values and all the ground truth values into z-scores: z-score(*x*) = (*x − µ*)*/σ*, where *µ* is the mean and *σ* is the standard deviation observed in the training dataset. The observed means and standard deviations were recorded to perform consistent z-score (and reverse z-score, for output values) transformations when validating/testing/applying the trained GNN. We trained the GNN to minimize the mean squared error (MSE) between predicted and ground truth values of z-score(CAD-goodness). We used Adam [25] optimizer with learning rate of 0.001.

We trained the GNN for 300 epochs. After every epoch we applied the trained GNN model to assess the best model selection rate using the validation dataset of interface graphs of docking models. We performed this training/validation process for multiple alternative GNN architectures. Figure S3 shows comparison of validation results (best model selection rates) for different hyperparameters: dropout (0.1 vs 0.25), number of attention heads (4 vs 8), number of hidden layers (3 vs 4). We then selected the best-performing GNN architecture (the one described in Figure 4). After choosing the GNN hyperparameters using the validation dataset, we verified that our choices were also beneficial for the performance on the testing dataset (Figure S4).

The training and validation process was repeated three times for the chosen architecture, giving us three selected GNN models (trained for 251, 89, and 297 epochs). In the VoroIF-GNN software the three trained GNN models are used as an ensemble — their output values are averaged.

### 2.8 Testing using docking models

We tested performance of multiple global scores for selecting best docking models using our prepared testing dataset. The testing results are summarized in Figure S5. Apart from the ideal selector (interface CAD-score computed using target structures), the best performing score is VoroIF-GNN-score. Most importantly, all the VoroIF-GNN scores significantly outperform our previous best interface-focused scoring method, VoroMQA-energy [26].

Interestingly, both VoroIF-GNN-score and VoroMQA-energy perform worse when normalized (either divided by the total interface area or averaged per interface residue). However, the normalization-related performance drop is significantly smaller in the case of VoroIF-GNN scores.

The average interface CAD-score of the models selected by the ideal selector and by VoroIF-GNN-score differ only by 0.03 (0.78 vs 0.75) — this means that VoroIF-GNN often selected accurate models even when failing to recognize the absolute best model.

## 3 Results

### 3.1 Overview of VoroIF-GNN application in CASP15

For the CASP15 EMA experiment, we registered two groups that employed the VoroIF-GNN method (“VoroIF” and “Venclovas”). For comparison we also registered “VoroMQA-select-2020”, representing our EMA method used back in CASP14.

CASP15 EMA rules allowed to submit two types of estimated global accuracy scores for every oligomeric structural model. The *first score* was intended for the overall structure accuracy of an oligomeric model, whereas the *second score* was supposed to represent the overall accuracy of the inter-chain interface. The specific global scoring methods used to derive these two types of scores for each group are summarized in Table 2.

**Table 2:**
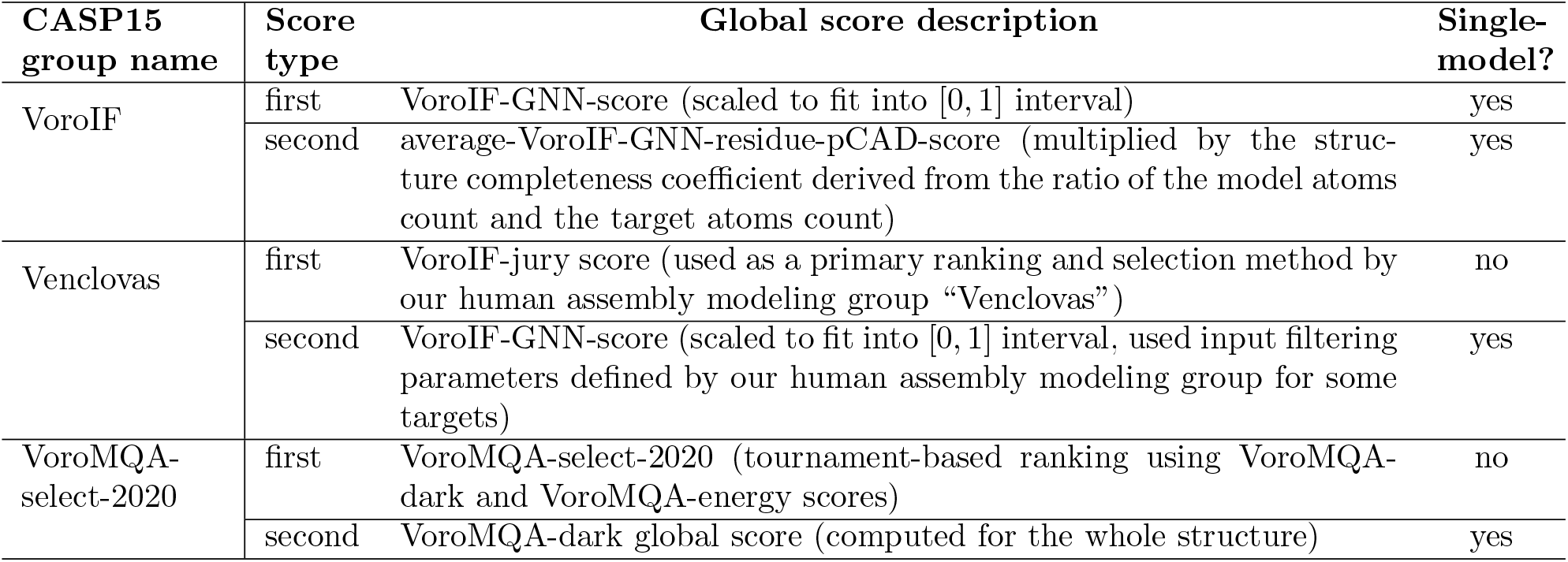
Global scoring methods behind our “VoroIF”, “Venclovas”, and “VoroMQA-select-2020” EMA groups in CASP15.

| CASP15 group name | Score type | Global score description | Single-model? |
| --- | --- | --- | --- |
| VoroIF | first | VoroIF-GNN-score (scaled to fit into $[0, 1]$ interval) | yes |
|  | second | average-VoroIF-GNN-residue-pCAD-score (multiplied by the structure completeness coefficient derived from the ratio of the model atoms count and the target atoms count) | yes |
| Venclovas | first | VoroIF-jury score (used as a primary ranking and selection method by our human assembly modeling group “Venclovas”) | no |
| | second | VoroIF-GNN-score (scaled to fit into $[0, 1]$ interval, used input filtering parameters defined by our human assembly modeling group for some targets) | yes |
| VoroMQA-select-2020 | first | VoroMQA-select-2020 (tournament-based ranking using VoroMQA-dark and VoroMQA-energy scores) | no |
|  | second | VoroMQA-dark global score (computed for the whole structure) | yes |

The default version of the method presented in this paper, VoroIF-GNN, was tested through “VoroIF” group. As explained in Table 2, for the first global score type we used VoroIF-GNN-score, for the second — average pCAD-score of the interface residues weighted by model completeness. VoroIF-GNN was also used by “Venclovas” group in two ways: for the second global score, and as a major component of the newly developed VoroIF-jury scoring method. VoroIF-jury is described in the accompanying publication dedicated to the methodology and results of the protein assembly modeling by the “Venclovas” group [11].

### 3.2 Summary of model selection performance

To evaluate the model selection performance, we first downloaded all the CASP15 EMA submissions from the CASP website. For every CASP multimeric target we selected the best models according to all the available global scores. For every global scoring method (of either first or second type) we calculated the sums of reference-based scores of the selected models. We chose to use score sums instead of score losses because not all groups submitted predictions for all the targets. We used four reference-based scores computed by the CASP15 organizers: ICS (F1-score) [27], IPS (Jaccard coefficient) [27], TM-score [28], and lDDT-oligo [29]. The first two scores measure the accuracy of the interface whereas the latter two assess overall structure accuracy. The ICS and TM-score sums are presented in Figure 5, separately for the first and the second global scores provided by CASP groups (some groups did not submit second or first scores). The IPS and lDDT-oligo sums are similarly presented in Figure S6. Overall, our methods performed relatively well. Based on the first score, the performance of “Venclovas” group was the best, but the underlying VoroIF-jury method is not a single-model method — it uses multiple models to compute final scores used for ranking. The best-performing single-model method, especially in terms of superposition-free accuracy (ICS, IPS, and lDDT-oligo), was VoroIF-GNN that was used by the “VoroIF” group for both the first and the second global scores.

**Figure 5:**
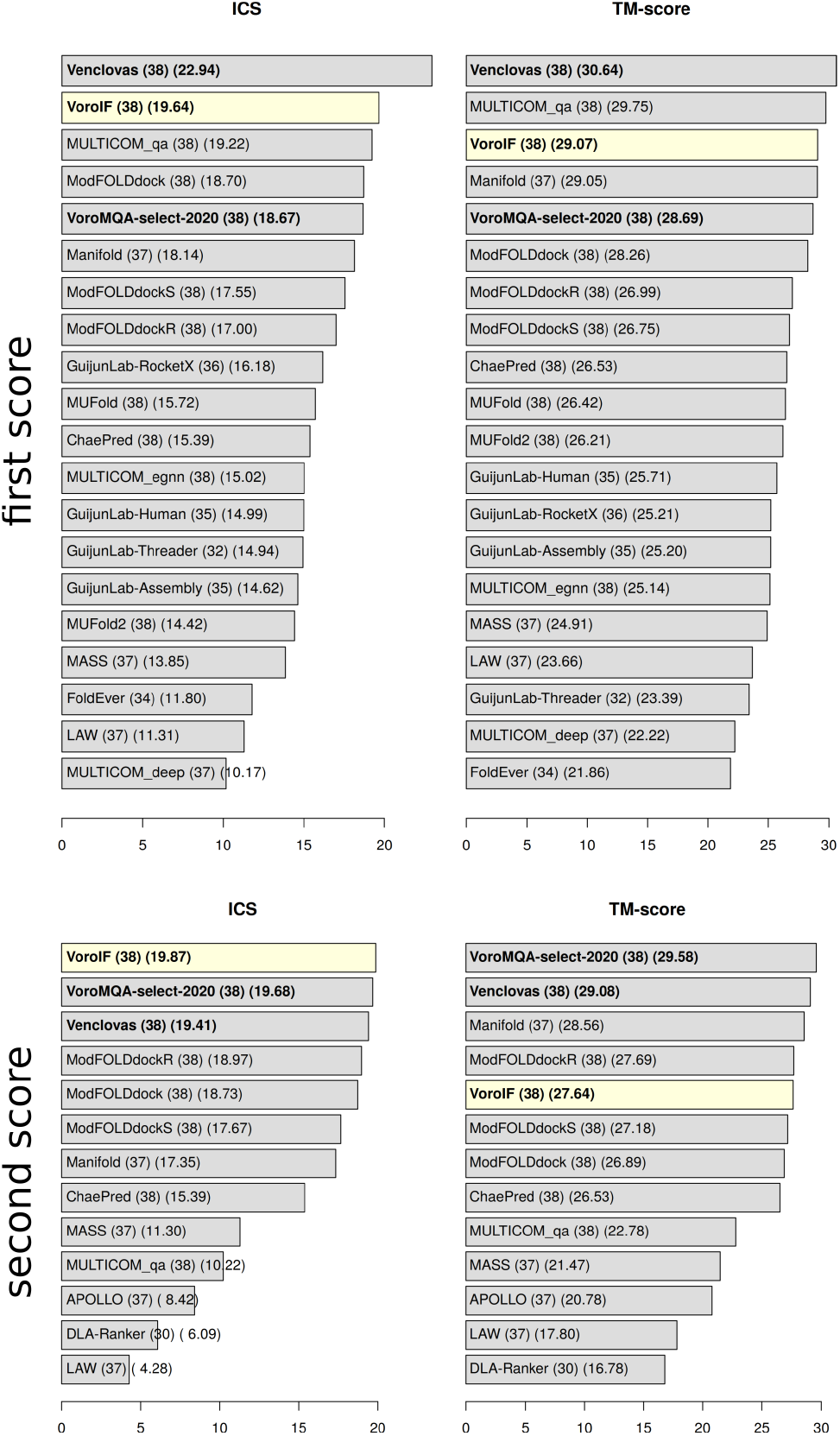
Sums of reference-based scores (ICS and TM-score) of first-ranked models for every CASP15 group. Label format: Group name (number of submissions) (sum of reference-based scores of selected models). Names of our groups are written on bold. “VoroIF” group, that relied solely on the VoroIF-GNN method, is highlighted in yellow.

### 3.3 Cases of failed interface selection

We looked at targets for which either the first (VoroIF-GNN-score) or the second (average-VoroIF-GNN-residue-pCAD-score) “VoroIF” group global score selected models with ICS further than 0.4 from the best available ICS. Such targets are highlighted in Figure S7. Both VoroIF-GNN global scores failed for targets H1129, H1142, T1160, and T1187. In addition, as highlighted in Figure S8, VoroIF-GNN-score failed for T1153, T1161, and T1179, while average-VoroIF-GNN-residue-pCAD-score failed for T1132, H1141, H1144, and T1170. Through visual analysis of the target and model structures we identified some factors that might have contributed to the VoroIF-GNN selection failures (briefly listed in Table S1).

In the case of H1129 (Suppl. Fig S9), one of the sequences provided for prediction was longer than in the solved target structure, and the extra tail in the model interfered with the interface. In this case the selection failure could have been avoided if our automated EMA pipeline included checking for clashes or topological knots.

In the cases of T1132 and T1170 (Suppl. Fig S10), VoroIF-GNN-score (the total summed predicted CAD-goodness of interface contacts) made relatively good selections, but average-VoroIF-GNN-residue-pCAD-score failed because it considered only the average predicted accuracy of contacts and disregarded the interface size.

In the cases of T1161 and T1179 (Suppl. Fig S11), some incorrect models had larger interface due to chain swapping that turned some intra-monomer contacts into inter-chain contacts. This resulted in bigger interfaces with some realistic contacts — such interfaces were interpreted overly favorably by VoroIF-GNN-score. Unlike the cases of T1132 and T1170, for T1161 and T1179 models the score normalization had a positive effect — average-VoroIF-GNN-residue-pCAD-score selected models with ICS *≥* 0.8.

In the case of the T1153 dimer (Suppl. Fig S12), the interface is relatively small considering the size of the monomers (297 residues). VoroIF-GNN-score was “fooled” by bigger interfaces among models, while average-VoroIF-GNN-residue-pCAD-score was not fooled because the average predicted accuracy of contacts in bigger interfaces was lower. However, we suspect that the primary reason for the failure of VoroIF-GNN-score may be a missing structural context. Using PPI3D [18] we found that T1153 is homologous to proteins that interact with DNA, e.g. bovine DNase I (PDB ID: 1dnk) shown in Suppl. Fig S12. If DNase I is superimposed with a relatively accurate dimeric model of T1153, the DNase I-bound dsDNA fits virtually perfectly inside the dimer. This observation suggests that in the absence of DNA the target structure of T1153 may not be biologically relevant.

In the case of the T1187 dimer (Suppl. Fig S13), the interface has a large empty cavity, which is an uncommon situation in biological assemblies. We could not find any information about the possible role of that cavity.

In the case of T1160, a small protein (48 a.a.), a significant part of the dimeric structure was not resolved. Therefore, the model scoring in this case was mostly influenced by the structural regions of the models with no counterparts in the experimentally solved structure. Moreover, according to the information provided by the CASP organizers, target structures of T1160 and T1161 (both very similar in sequence) might have been significantly influenced by crystallization conditions. VoroIF-GNN was not trained to account for such factors.

H1141, H1142, and H1144 represent three of the five complexes (H1140-H1144), in which the same protein (mouse CNPase) is bound to different nanobodies. It is a common knowledge that nanobody and antibody binding assemblies are often difficult to predict and score. In the case of H1142, there were no highly accurate models to select from (best available ICS= 0.447). In the cases of H1141 and H1144, VoroIF-GNN-score selected good models (with ICS of 0.75 and 0.73, respectively), but average-VoroIF-GNN-residue-pCAD-score failed, once again indicating that normalizing predicted accuracy by interface size is not always beneficial.

It is also worth noting that unnormalized VoroIF-GNN-score performed better in terms of the total sum of TM-scores of selections for all CASP15 targets (Figure 5). Nevertheless, keeping in mind the cases of possible lack of context (T1153) or significant differences among modeled monomers resulting in over-prediction of the interface area (T1161 and T1179), we cannot conclude that the unnormalized global VoroIF-GNN score is always better than the normalized one. It may be advantageous to use both to get a better characterization of a modeled interface.

## 4 Conclusions

VoroIF-GNN is first and foremost a proof-of-concept method. We developed this method to test the idea of deriving protein-protein interface graphs using Voronoi tessellation and to explore the suitability of such graphs for training graph neural networks. CASP15 results showed that VoroIF-GNN is a robust method for estimating accuracy of protein assemblies, in particular when judged by superposition-free measures of interface and structure accuracy. For practical applications, VoroIF-GNN has several advantages: it only requires a single structure of protein complex for input; it does not depend on the structure prediction method that was used to derive a protein complex; it works for protein complexes with any stoichiometry and for any number of inter-chain interfaces. As for limitations, VoroIF-GNN is not an all-encompassing method, as it does not try to assess all the properties of a protein-protein complex. However, due to novelty of the VoroIF-GNN methodology, VoroIF-GNN scores can be especially informative and useful when combined with other scores in a meta-method as exemplified by VoroIF-jury.

## Availability

VoroIF-GNN software is freely available as part of Voronota-JS at https://kliment-olechnovic.github.io/voronota/expansion_js/.

Docking models used to train, validate, and test VoroIF-GNN are available at https://dx.doi.org/10.5281/zenodo.7841307.

All our CASP15 pipeline scripts are available as part of FTDMP at https://klimentolechnovic.github.io/ftdmp/.

## Acknowledgments

We thank Justas Dapkūnas and Simonas Vytautas Ašmontas for valuable comments about the manuscript.

