## Supplementary information for "VoroIF-GNN: Voronoi tessellation-derived protein-protein interface assessment using a graph neural network"

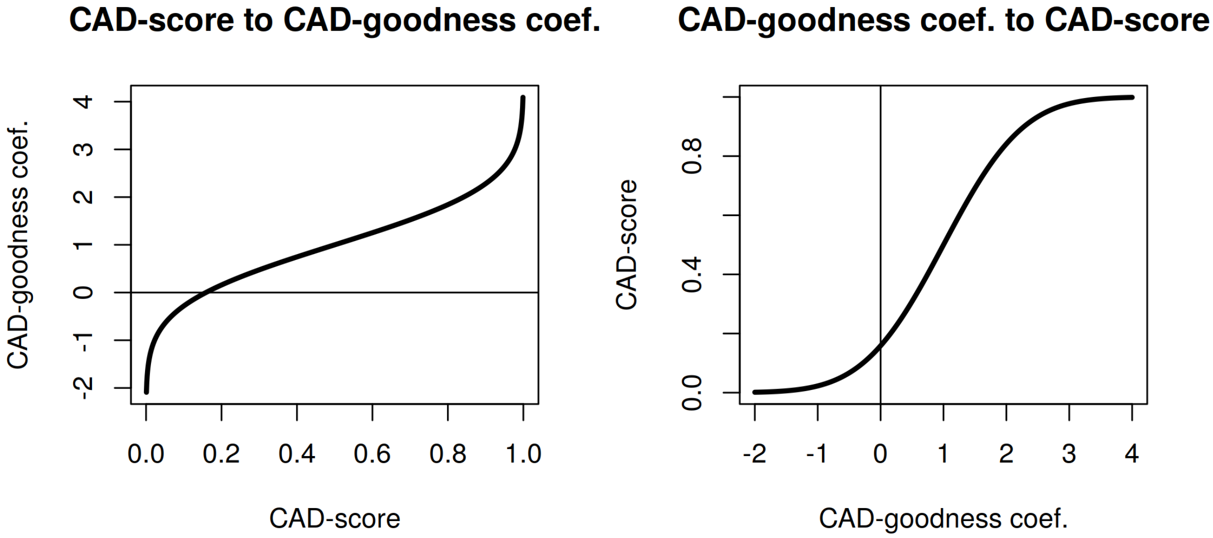


**Figure S1:** Converting contact CAD-score value to CAD-goodness coefficients and back.

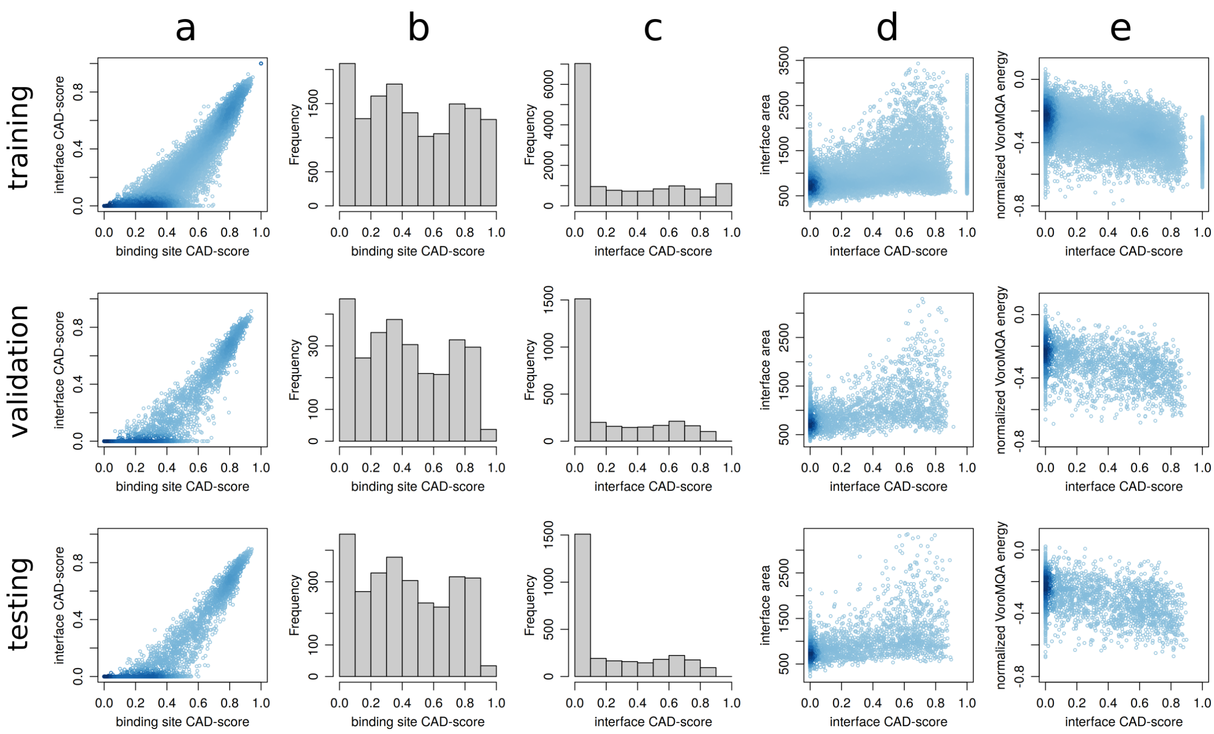


**Figure S2:** Summary of binding site CAD-score and interface CAD-score values of the models in the training, validation, and testing datasets: relations between interface and binding site CAD-scores (a), histograms of interface CAD-scores (b) and binding site CAD-scores (c), relations between interface CAD-score and interface area (d), relations between interface CAD-score and normalized VoroMQA energy (e). Scatter plots are colored according to point density.

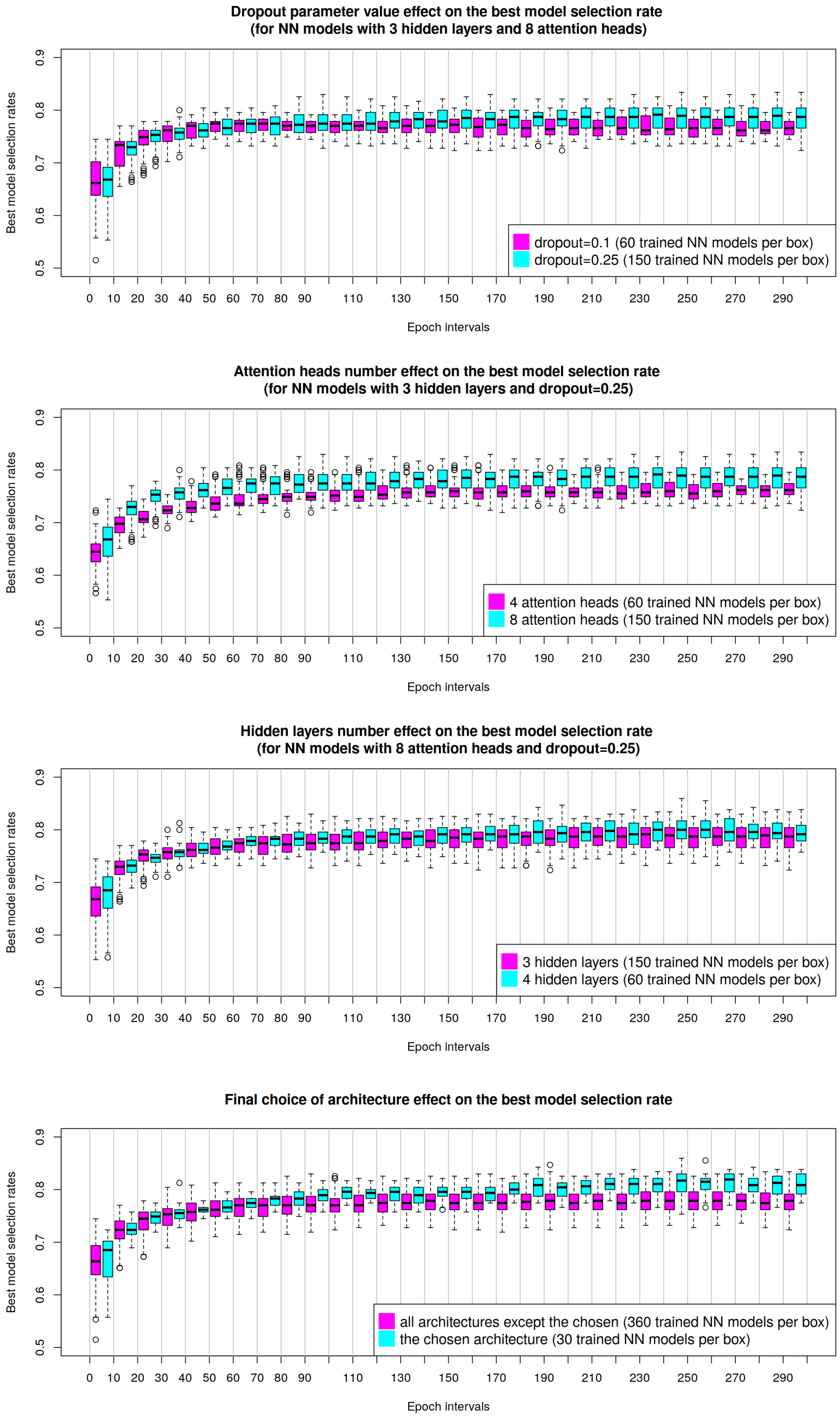


**Figure S3:** Validation set statistics of best model selection rates (for intervals of 10 training epochs) depending on various neural network hyperparameters.

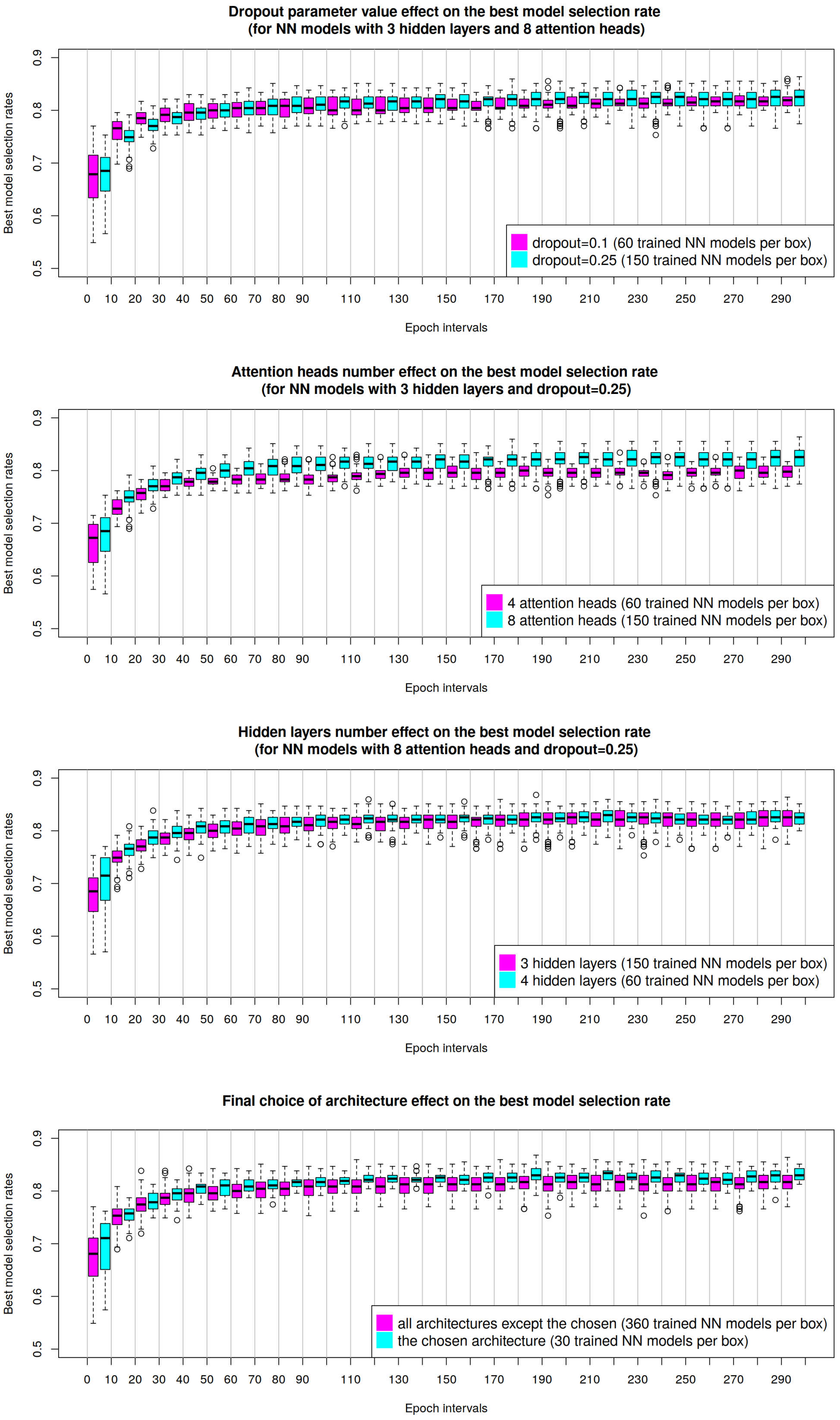


**Figure S4:** Testing set statistics of best model selection rates (for intervals of 10 training epochs) depending on various neural network hyperparameters.

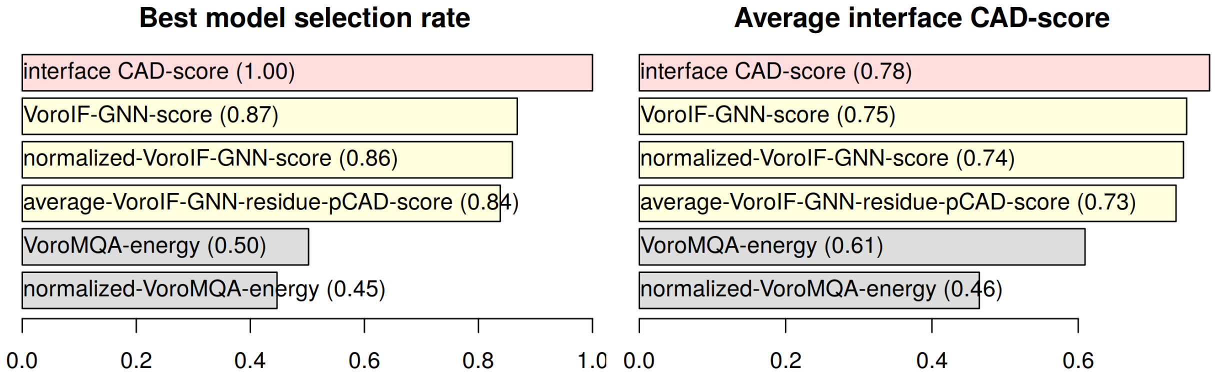


**Figure S5:** Docking model selection performance for the ideal selection score (red), VoroIF-GNN scores (yellow), and VoroMQA-energy scores (gray). The testing was done using 235 sets of models.

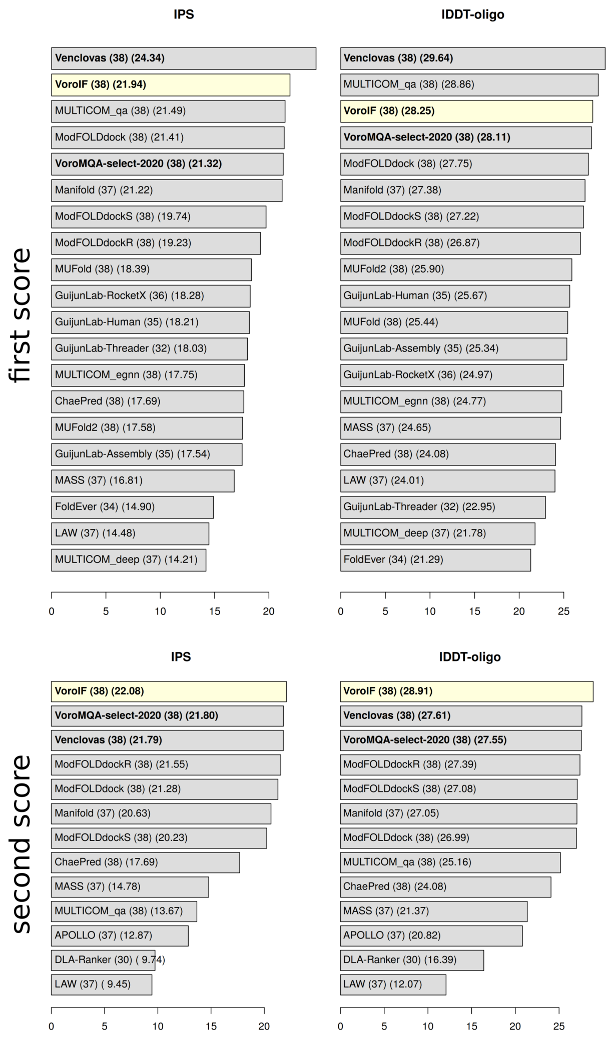


**Figure S6:** Sums of reference-based scores (IPS andlDDT-oligo) of first-ranked models for every CASP15 group. Label format: Group name (number of submissions) (sum of reference-based scores of selected models). Names of our groups are written on bold. “VoroIF” group, that relied solely on the VoroIF-GNN method, is highlighted in yellow.

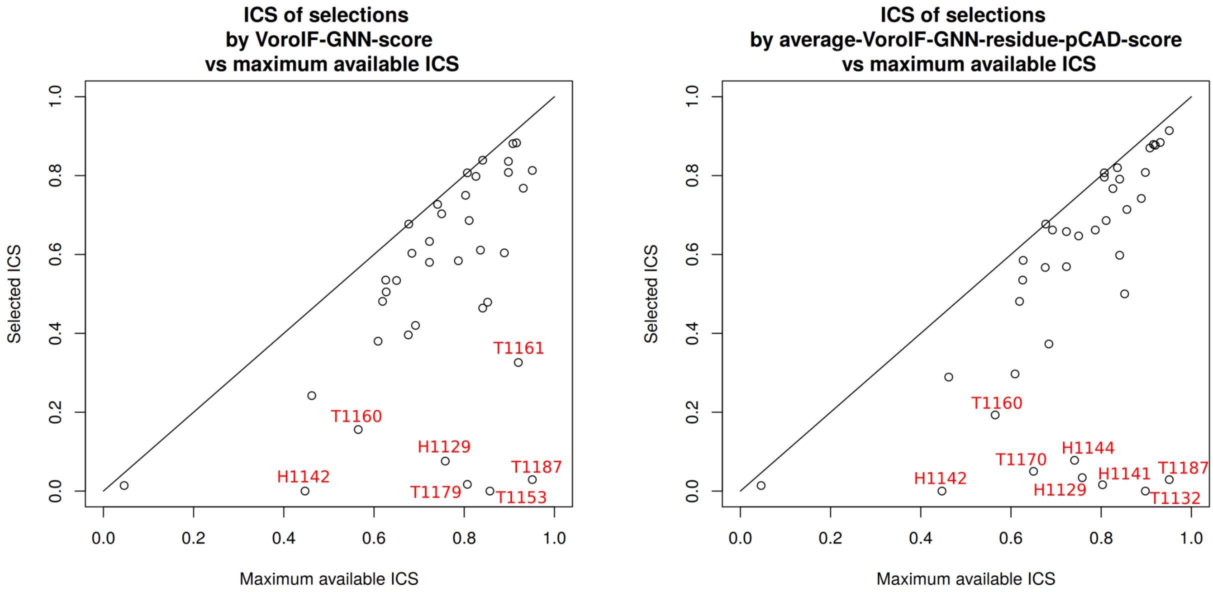


**Figure S7:** Per-target performance of VoroIF-GNN global scoring in CASP15. For every target, ICS of the selected model is plotted against the maximum achieved ICS for that target.

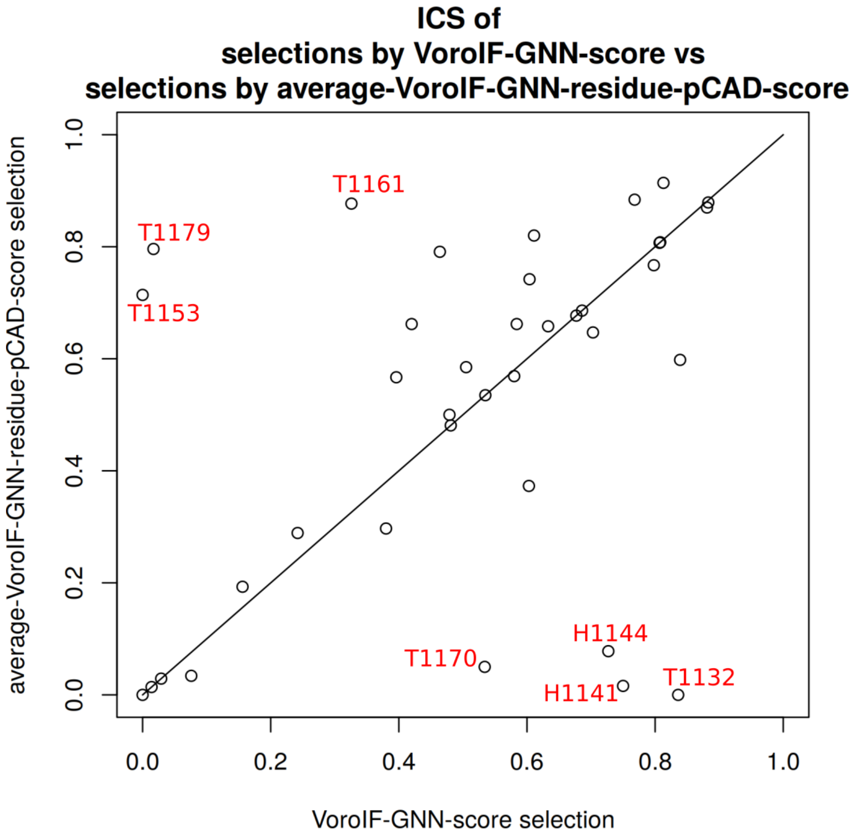


**Figure S8:** Per-target comparison of model selections made by different VoroIF-GNN global scores in CASP15.

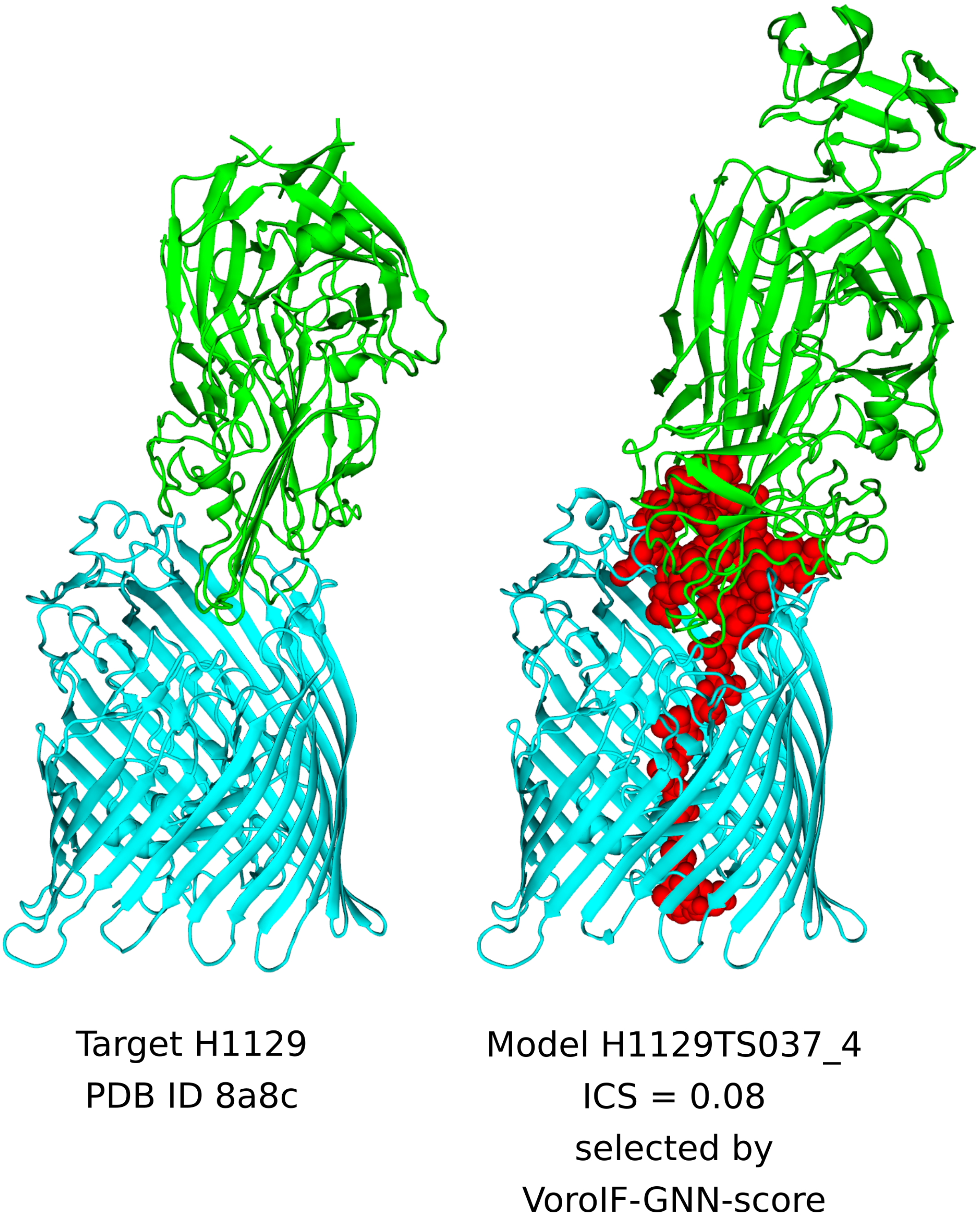


**Figure S9:** Model selection failure case study for target H1129.

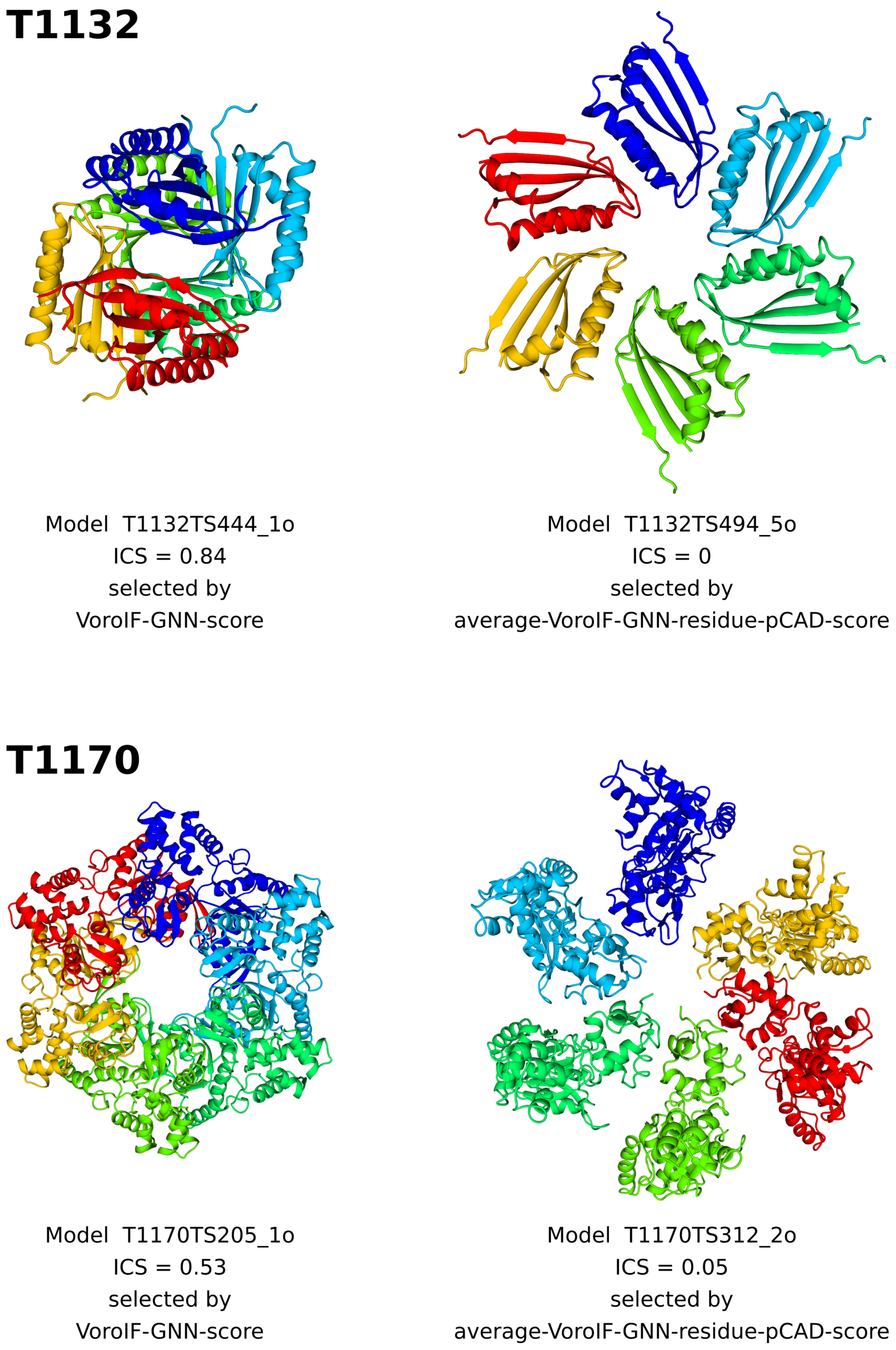


**Figure S10:** Model selection failure case studies for targets T1132 and T1170.

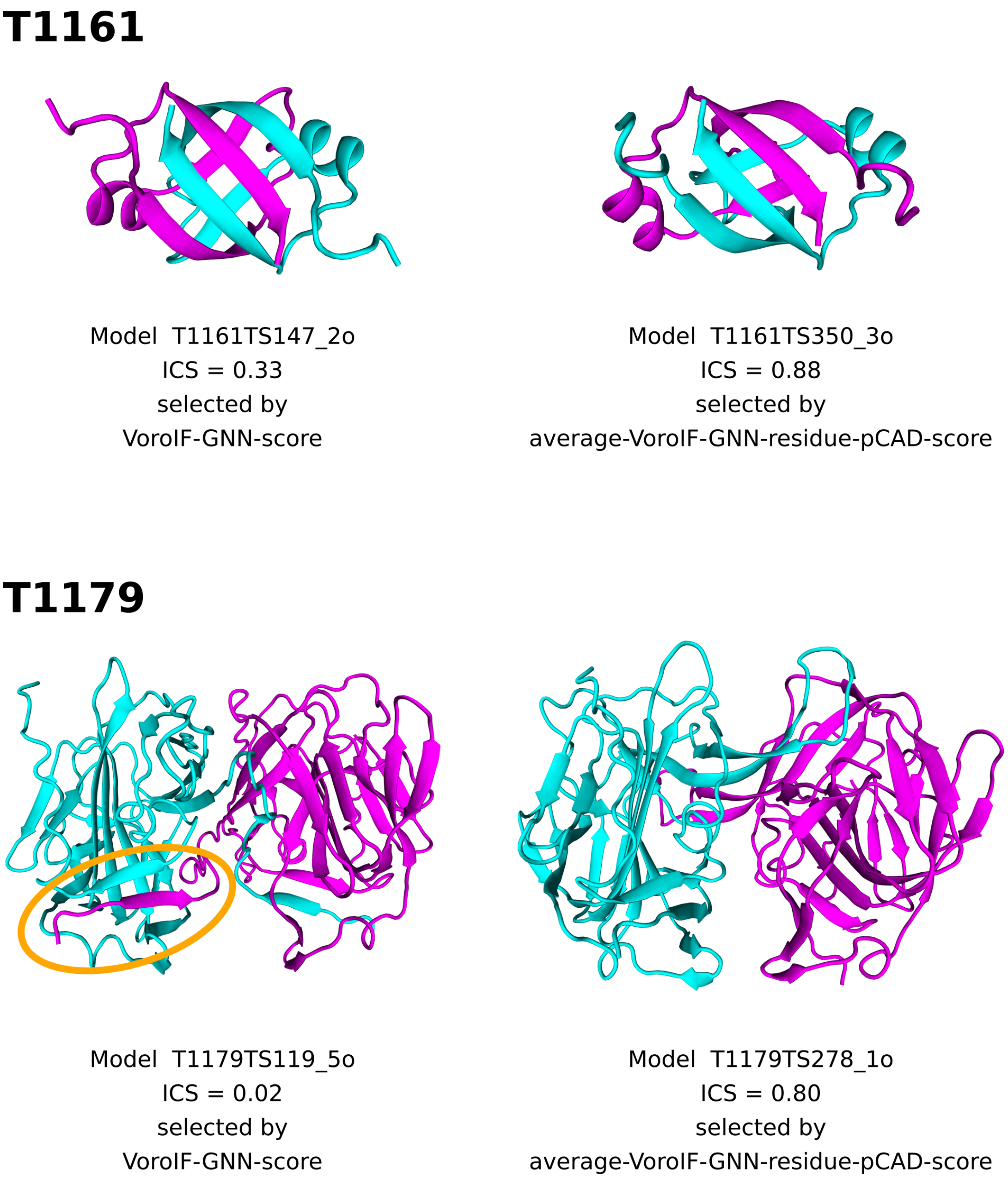


**Figure S11:** Model selection failure case studies for targets T1161 and T1179.

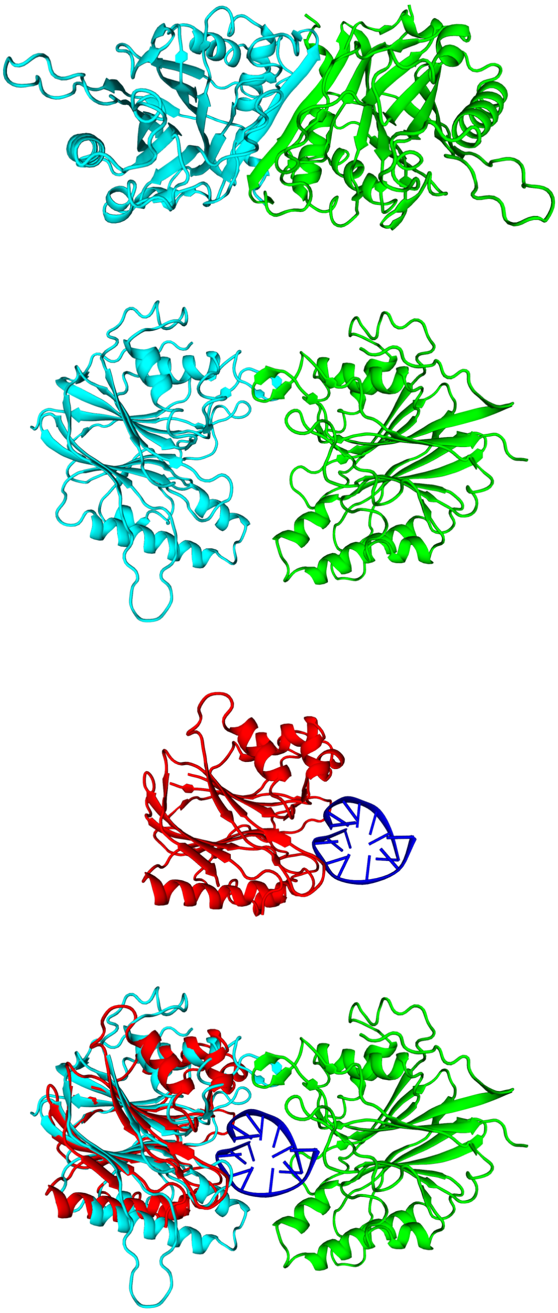

Model T1153TS278_4o
ICS = 0
selected by
VoroIF-GNN-score

Model T1153TS185_3o
ICS = 0.71
selected by
average-VoroIF-GNN-residue-pCAD-score

Homologous
protein-DNA complex
PDB ID 1dnk

1dnk
aligned on
T1153TS185_3o

**Figure S12:** Model selection failure case study for target T1153.

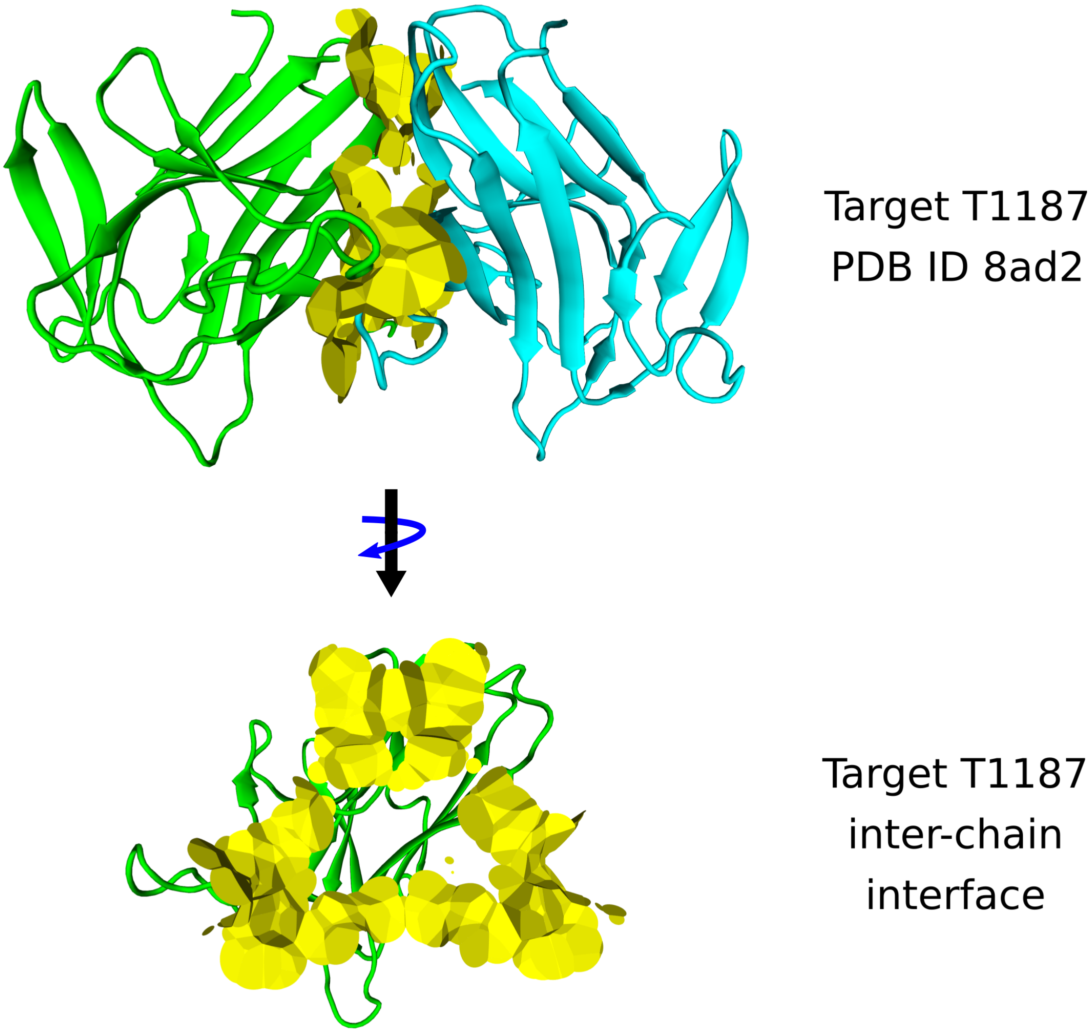


**Figure S13:** Interface of target T1187.

**Table S1:** Factors that might have contributed to the VoroIF-GNN selection failures.

| | |
|---|---|
| **Both VoroIF-GNN scores failed** | |
| H1129 | Models have extra tails absent in the solved structure |
| H1142 | Nanobody complex (no highly accurate models) |
| T1160 | Models have extra tails; crystallization conditions? |
| T1187 | Large cavity within the interface |
| **Only VoroIF-GNN-score failed** | |
| T1153 | Missing bound DNA? |
| T1161 | Non-swapped vs. swapped chains; crystallization conditions? |
| T1179 | Non-swapped vs. swapped chains |
| **Only average-VoroIF-GNN-residue-pCAD-score failed** | |
| T1132 | Interface size |
| H1141 | Nanobody complex |
| H1144 | Nanobody complex |
| T1170 | Interface size |
